# Identifying Distinct Molecular Response of CAR-T cells to Solid Tumors by Synthetic Single-Cell Transcriptomic Analyses

**DOI:** 10.1101/2025.09.17.676755

**Authors:** Yi Hui Luo, Elisabetta Forte, Yoshiaki Tanaka

## Abstract

Chimeric Antigen Receptor (CAR)-T cell therapy is a novel personalized treatment that engineers patient immune cells to fight against cancer cells. Recently, CAR-T cell therapy has demonstrated remarkable success in the treatment of hematopoietic cancers, whereas the treatment of solid tumors is more challenging, likely in part due to their severely immunosuppressive tumor microenvironment. To address distinct molecular responses of CAR-T cells between blood and solid tumors, we performed synthetic analysis of single-cell transcriptomics of CAR-T cells and identified unique immunosuppressive subpopulations and aberrant signalling inductions in CD4^+^ and CD8^+^ CAR-T cells in the context of solid tumors. Furthermore, we also found that PD-1-independent exhaustion-like CD8^+^ CAR-T cells, characterized by high expression of *TNFRSF9* and *CCL3*, were preferentially generated under solid tumor stimulation. Collectively, our comprehensive analyses provide essential molecular insights into solid tumor-stimulated CAR-T cells and assists in overcoming the limited efficacy of CAR-T cell therapy against solid tumors.

## Introduction

Chimeric Antigen Receptor (CAR) is a recombinant receptor that can ignite antigen targeting and cytotoxicity functions into T cells [1]. CAR-T cells infused into the patients have persistent anti-tumor activity, and lead to highly durable remission. Impressively, CAR-T cell therapies have demonstrated remarkable efficacy in the treatment of hematopoietic cancers [2]. So far, Food and Drug Administration (FDA) has approved five CAR-T cell products (Kymriah, Yescarta, Tecartus, Breyanzi, and Aucatzy) that targets the CD19 antigen and are used in the treatment of precursor B-cell acute lymphoblastic leukemia (B-ALL) or diffuse large B-cell lymphoma (DLBCL) [3]. The other two FDA-approved CAR-T cells (Abecma and Carvykti) bind BCMA, whose expression is generally elevated in patients with multiple myeloma (MM).

CAR construct is one of major determinant for the functionality of CAR-T cells and has been optimized by replacing domains. CAR is composed of four major domains: i) antigen-binding, ii) hinge, iii) transmembrane, and iv) intracellular signaling domain [4]. The antigen-binding domain usually forms a single-chain variable fragment (scFv) that is derived from the variable heavy (VH) and light chain (VL) of immunoglobulin G (IgG). The hinge (also known as spacer) domain is an extracellular structural region of CAR connecting the antigen-binding and the transmembrane domain, and essential for spatial flexibility between the antigen-binding domain and epitope, and stable expansion of T cells [5]. Although IgG hinge was originally used in this domain, CH2 region in IgG hinge is unintentionally targeted by innate immune cells, such as natural killer cells, and leads to cell death and rapidly loses the anti-tumor effect of CAR-T cells [6]. Thus, the hinge domain is now designed by IgG without CH2 region or non-IgG-based spacers, such as CD8α, CD28, and NGFR. The transmembrane domain serves as an anchor of the CAR structure on T cell membrane and propagates the antigen recognition signal into the intracellular domain [4]. This domain is typically designed from immune cell surface proteins, such as CD8α and CD28. The intracellular signaling domain is essential for activation of downstream signaling pathways, and further subdivided into co-stimulatory and activation domains [7]. Most of CARs employ immunoreceptor tyrosine-based activation motif of CD3ζ for the activation domain, whereas co-stimulatory domain is derived from CD28, 4-1BB (CD137), or OX40 (CD134), and others.

Despite recent successes in blood cancer treatment and the improvement of CAR constructs, CAR-T cell therapies still face pivotal challenges in the treatment of solid tumors, such as lung and brain cancers [4, 8]. Although several clinical trials against solid tumors are still underway, most of the disclosed trials could not obtain desirable outcomes [9]. For example, a recent phase I trial showed that CAR-T cells were infiltrated into glioblastoma multiforme (GBM) without any toxicities and side effects, but undesirably inactivated with elevation of regulatory and exhausted T cell markers [10]. Another phase I trial for non-small lung cancer also demonstrated substantial infiltration of CAR-T cells into tumors, but showed poor persistency [11]. A major obstacle is the immunosuppressive tumor microenvironment of solid tumors that is embedded with surrounding normal cells and leads to exhaustion of CAR-T cells [8]. Although many efforts have struggled with these issues by co-administrating CAR-T cell therapies with immune checkpoint inhibitors (e.g. anti-PD-1) or designing CARs targeting multiple antigens [4, 12], there is no FDA-approved CAR-T cell therapy against solid malignancies yet.

Gene expression profiling is a powerful tool to address how CAR-T cells are activated by exposure to tumors and dampened by their immunosuppressive microenvironment. Recent preclinical and clinical studies have occasionally employed single-cell RNA-sequencing (scRNA-seq) of CAR-T cells to investigate their molecular responses against cancers, and deposited their data into public registries. Importantly, these public datasets were obtained under various conditions (e.g. blood or solid tumor, patients, animals, or *in vitro* stimulation). They are precious resources to robustly understand how CAR-T cells respond to cancer in different manners across tumor types, CAR constructs, and experimental models. Here, we systematically collected public CAR-T cell scRNA-seq datasets, and demonstrated how CAR-T cell transcriptomics are altered across cancer types, experimental models, and CAR structures.

## Results

### Organization of the collected single-cell transcriptomics

Throughout public resources, total 362 scRNA-seq libraries of CAR-induced and control non-induced T cells from 40 different projects were collected (See data collection rules in Materials and Methods) (Figure 1A, Table S1). Sample information, such as target cancer type, CAR construct, and experimental models, was manually collected from literature or meta-data tables in public repositories (Figure 1B). 157 and 58 CAR-induced T cell libraries were against blood and solid tumors, respectively, whereas 101 were without any tumor stimulation (Figure 1C). 46 libraries were for non-induced T cells (T cells before CAR induction or CAR^-^ fraction). The blood cancer data were mostly from FDA-approved targets (e.g. MM, B-ALL, DLBCL), but some of them were from non-approved cancer types (e.g. Plasma cell leukemia (PCL)). The solid tumor data included brain, pancreas, lung, bone, and prostate tumors. Around half of the collected datasets were derived from patients, and the others were obtained by *in vitro* co-culture with cancer cells or from mice bearing tumor xenografts (Figure 1D). Solid tumor-stimulated CAR-T cells from human and xenograft studies were isolated from resected tumors and are therefore tumor-infiltrating lymphocytes. The collected datasets contain CARs with scFv for 14 different target antigens (e.g. CD19 and BCMA), 5 hinge (e.g. CD8α and IgG), 3 transmembrane (e.g. CD8α, CD28), and 4 co-stimulatory domains (e.g. 4-1BB, CD28) (Figure 1E). Although almost all CARs used CD3ζ-based activation domain, ZAP70-based activation domain was also used in a few studies. After quality control and batch effect corrections, about 1.2 million individual cells were separated into CD4^+^ and CD8^+^ T cells (Figure S1A-C). Our relatively stringent cutoffs excluded 28.9%±12.2% of cells in the quality control step in each dataset (Figure S1D). In addition, 1.70%±1.51% of cells were categorized into CD4 and CD8 double positive T cells. Recent flow cytometry analyses implied that CD4 and CD8 double positive T cells substantially exist in peripheral blood of the tumor patients [13]. However, we could not rule out the possibility of artificial doublets for these double positive cells (Figure S1E). Therefore, we used 411,136 and 845,357 single CD4^+^ and CD8^+^ T cells for subsequent analyses, respectively.

**Figure 1.**
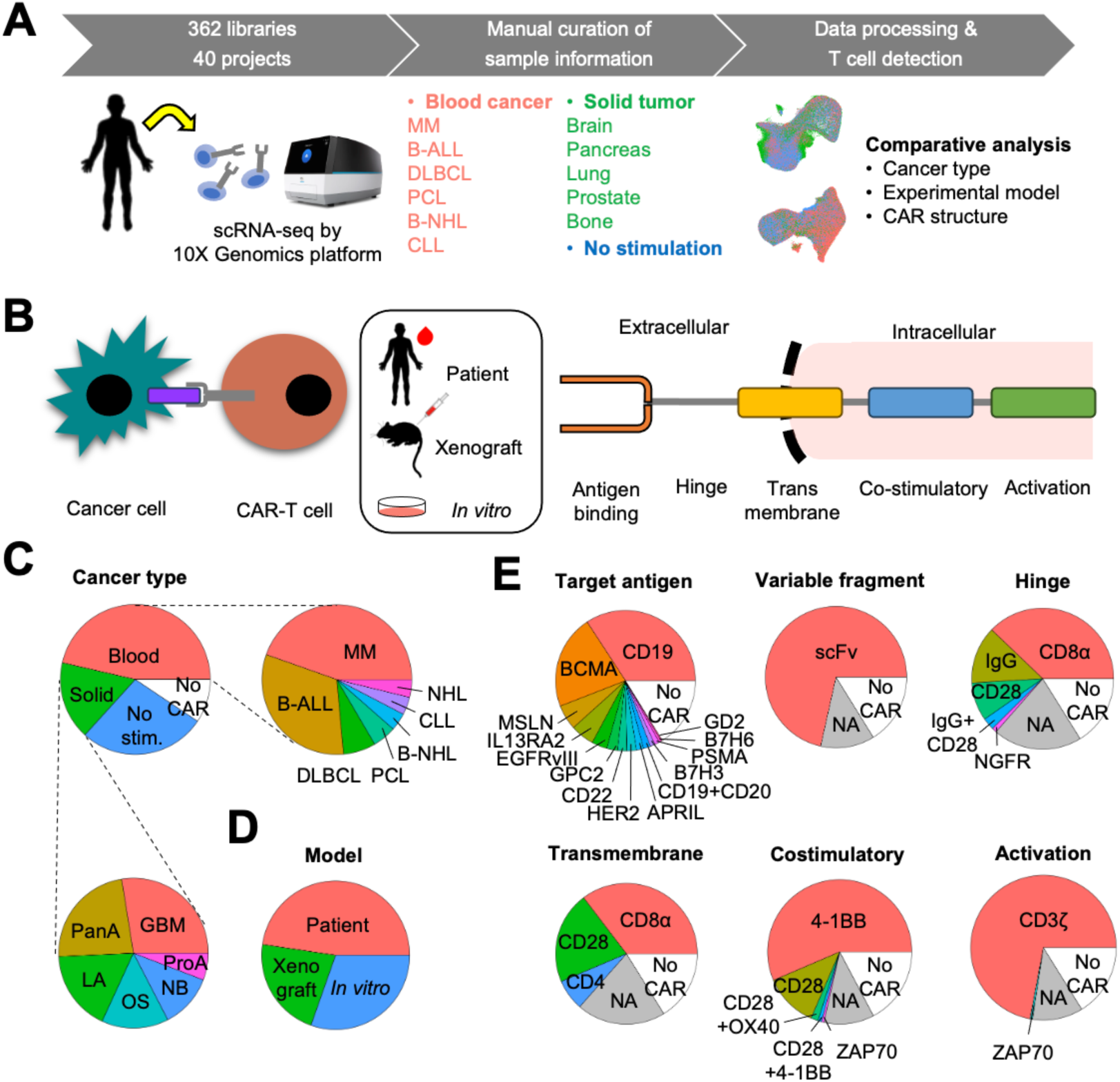
Comprehensive collections of single-cell transcriptomics of CAR-T cells. **A.** Analytic workflow of comprehensive analysis of single-cell transcriptomcis of CAR-T cells. **B.** Schematics of classification of scRNA-seq of CAR-T cells. The single-cell transcriptomics were classified by experimental models, target cancer types, and CAR constructs. **C-E.** Pie charts showing statistics of the collected scRNA-seq libraries Cancer type (**C**), experimental model (**D**), and CAR strcutures (**E**) are shown. No stim.: no tumor-stimulation sample, No CAR: no CAR induction, MM: multiple myeloma, B-ALL: B Cell acute lymphoblastic leukemia, DLBCL: diffuse large B cell lymphoma, PCL: plasma cell leukemia, B-NHL: B-cell non-Hodgkin’s lymphoma, CLL: chronic lymphocytic leukemia, NHL: non-Hodgkin’s lymphoma, CML: chronic myelogenous leukemia, GBM: glioblastoma multiforme, PanA: pancreatic adenocarcinoma, LA: lung adenocarcinoma, OS: osteoblastoma, NB: neuroblastoma, ProA: prostate adenocarcinoma

### Identification of solid tumor-specific subpopulation in CD4^+^ CAR-T cells

CD4^+^ T cells play important roles in antitumor immunity by stimulating other immune cells and coordinating their activity. After data preprocessing, the collected CD4^+^ T cells were clearly segregated by *CD3* family genes that are essential receptors for T cell activation (Figure 2A and S2A). *CD4* is a major T helper cell surface marker, but also expressed in human monocyte [14]. Since monocytic markers, such as *CD14* and *CD16*, were highly expressed in CD3^-^ fraction (Figure S2A), we interpreted CD3^-^ fraction as CD4^+^ monocytes. We observed that CD4^+^CD3^+^ T cells were predominant in CAR-induced scRNA-seq datasets with pre-sorting of CAR construct (Figure 2B and Table S1). Furthermore, a portion of CD4^+^CD3^+^ T cells were detected almost exclusively in CAR-induced datasets (defined as “CD3^+^ unique”) and characterized by high expression of cell cycle-related genes (Figure 2C). In contrast, genes involved in cytokine production, such as tumor necrosis factor (TNF) and interleukin (IL) 1β, 6, and 10, were significantly up-regulated in CD4^+^ monocytes (Figure 2C). These cytokines are known to be associated with cytokine release syndrome, which is a common side effect of CAR-T cell therapy [15-18]. We found that these cytokine-encoding genes (e.g. *IL1B*, *TNFSF13*) were highly expressed in CD4^+^ monocytes but mostly diminished in CD4^+^CD3^+^ T cells (Figure 2D). *PFKFB4*, which is a potential prognostic marker of cytokine release syndrome grade [19], was also highly expressed in CD4^+^ monocytes. Furthermore, *PFKFB4* expression in CD4^+^ monocytes was significantly increased post infusion of CAR-T cells (Figure S2B, p<3.96e-2 by two-sided T test). These results are consistent with a previous finding that cytokine release syndrome is mainly mediated by monocyte-derived cytokines, but not CAR-T cell [20].

**Figure 2.**
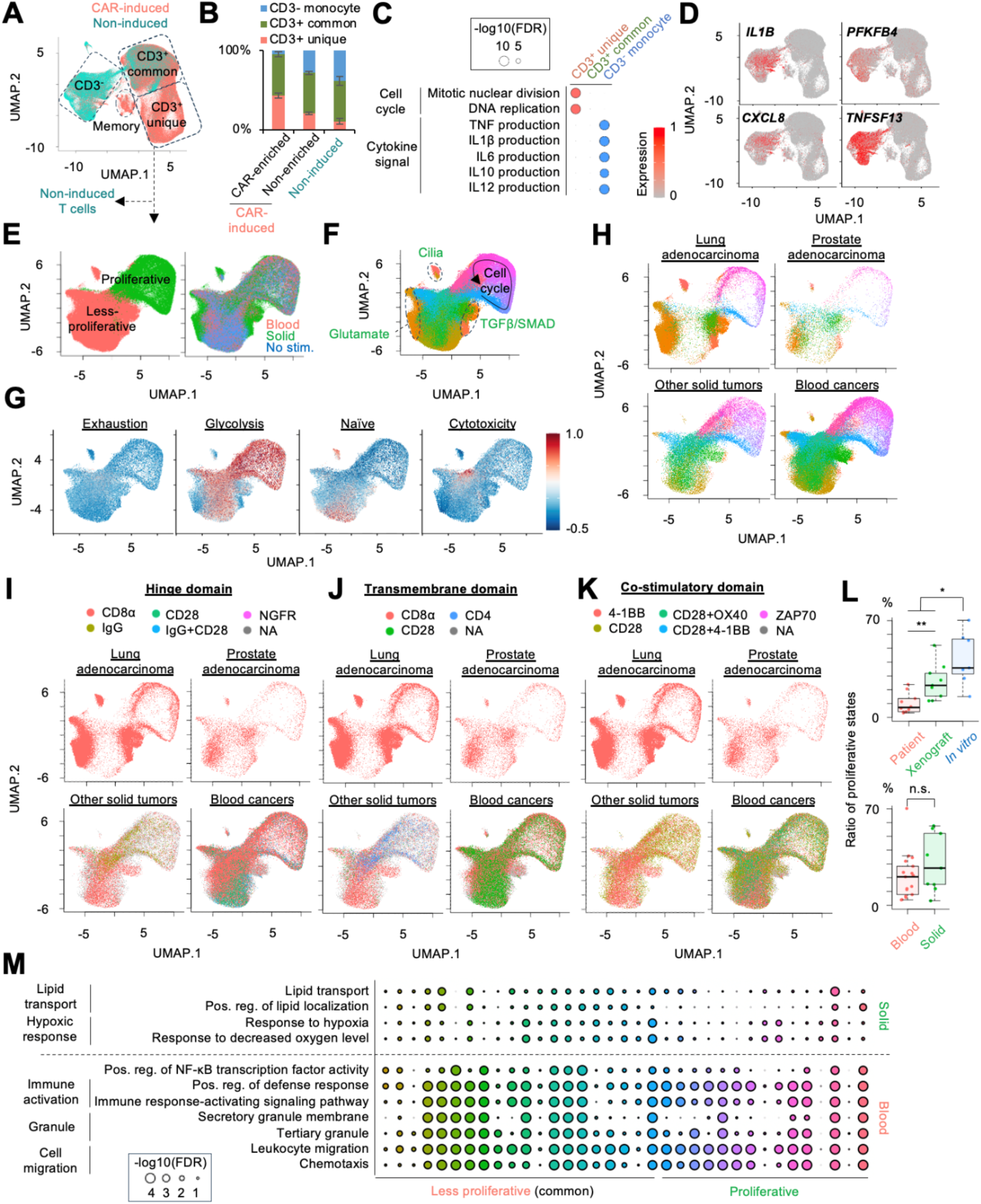
Identification of unique characteristics in blood and solid tumor-stimulated CD4^+^ CAR-T cells. **A.** UMAP plot of individual CD4^+^ T cells from 40 different projects. CAR- and non-induced T cells were distinguished by different colors. **B.** Enrichment of CD4^+^ T cell fraction in CAR- and non-induced T cell datasets. **C.** Representative significant GO terms between CAR- and non-induced CD4^+^ T cells. **D.** Feature plot representing expression of cytokine release syndrome-related genes in CD4^+^ T cells. **E.** UMAP plot of individual CD4^+^ CAR-T cells colored by proliferative states and cancer stimulation. **F.** UMAP plot of individual CD4^+^ CAR-T cells colored by clusters. Representative annotations are also shown. Annotations for solid tumor-specific clusters are shown by green text. **G.** Feature plot representing enrichment of CD4^+^ T cell gene signatures. **H.** UMAP plot of individual CD4^+^ CAR-T cells distinguished by target cancer types. Colors correspond to clusters in Figure 2E. **I-K.** UMAP plot of individual CD4^+^ CAR-T cells distinguished by types of (**I**) hinge, (**J**) transmembrane, and (**K**) co-stimulatory domain. **L.** Comparison of the percentage of cells in proliferative state across experimental models and the target cancer types. * p<0.05 ** p<0.01 by two-sided T test **M.** Representative GO terms for differentially expressed genes between blood and solid tumor-stimulated CD4^+^ CAR-T cells in each common cluster. Circle size represent -log10(FDR).

To further dissect the molecular characteristics of CD4^+^ CAR-T cells, we performed re-normalization of CD4^+^CD3^+^ cell clusters after excluding non-induced datasets. Since specific CD4^+^ CAR-T cell clusters were predominantly derived from one project (SRP489891) [21] (Figure S2C), we excluded them from subsequent analyses. The re-normalization segregated CD4^+^ CAR-T cell clusters into two major states: proliferative and less proliferative state (Figure 2E and S2D). Intriguingly, certain less proliferative clusters were preferentially detected in CD4^+^ CAR-T cells stimulated by solid tumors, while proliferative clusters were universally detected in blood cancer-, solid tumor-, and non-stimulated CD4^+^ CAR-T cells (right panel in Figure 2E). Gene Ontology (GO) analysis for differentially expressed genes across clusters revealed that the solid tumor-specific less proliferative clusters were characterized by high expression of genes related to cilia, glutamate, SMAD and TGFβ signaling pathways (Figure 2F and S2E). Notably, glutamate, SMAD and TGFβ signals are involved in suppression of antitumor immunity and are associated with poor prognosis [22-24]. Furthermore, gene signature analysis for T cell function demonstrated that the solid tumor-specific clusters significantly down-regulated glycolysis pathways (Figure 2G) [25]. Interestingly, glycolysis level of CD4^+^ T cells is positively associated with their inflammatory effect and antitumor response, and negatively with their immunosuppressive functions [26, 27]. Therefore, these solid tumor-specific clusters were likely to harbor immunosuppressive characteristics. Although further investigation is still needed, our results imply that the derivation of immunosuppressive subpopulations may be associated with the limited efficacy of CAR-T cell therapy against solid tumors.

### Distinct subpopulation composition of blood cancer-stimulated CD4^+^ CAR-T cells between CD8α and CD28 hinge and transmembrane domain

Given the existence of the solid tumor-specific clusters, we next asked whether this CD4^+^

CAR-T cell subpopulation is associated with specific tumor types, CAR constructs, and experimental models. Surprisingly, the solid tumor-specific clusters were mainly detected in lung and prostate tumors, whereas other solid tumors displayed similar cluster composition as blood cancers (Figure 2H). In addition, we found that the subpopulation composition varies by CAR construct, particularly in blood cancer (Figure 2I-K). For example, the less-proliferative clusters characterized by enrichment of naïve T cell signatures (Figure 2G) consisted mostly of CD4^+^ CAR-T cells with CD28-hinge domain (Figure 2I). Three major CD28-hinge CAR-T cell datasets (>2,000 cells, SRP310699, SRP441576, and SRP475734) made up about 8-13% of the naïve T cell cluster [28-30], whereas 3-7% was from CD8α-hinge CAR-T cell datasets (SRP388806, SRP250485, and SRP470067) (Figure S2F) [31-33]. Although CD4^+^ CAR-T cells with 4-1BB co-stimulatory domain also seem to preferentially appear in the naïve T cell cluster (Figure 2K), this might be because two major CD28-hinge CAR-T cell datasets (SRP441576 and SRP475734) utilized 4-1BB co-stimulatory domain (Figure S2F). In addition to the hinge domain, the transmembrane domain affected the subpopulation composition of CD4^+^ CAR-T cells (Figure 2J). Especially, CD4^+^ CAR-T cells with CD8α transmembrane domain generated a substantial number of cells in SMAD/TGFβ signal-enriched clusters (0.14-0.28%), whereas no or very few cells were detected with CD28 transmembrane domain (0.00-0.02%) (Figure S2F). Importantly, recent studies demonstrated that CAR-T cells with CD28 hinge and transmembrane domain were more sensitive to low antigen concentrations, but were more vulnerable to activation-induced cell death than those with CD8α [34, 35]. In fact, the CD28-hinge domain-specific clusters highly expressed genes related to T cell activation (e.g. *AQP3*, *LINC00861, LEF1, TCF7*) (Figure S2G) [36-38]. These results suggest that the design of CAR construct potentially affects the derivation of specific CAR-T cell subpopulations, and our scRNA-seq collection is a precious resource to unravel the nature of the best CAR design for each tumor type.

### Reduction of proliferative CD4^+^ CAR-T cells in *in vivo* experimental models

According to the cluster distribution, IgG hinge and CD4 transmembrane domain also seem to be related to the composition of proliferative and less proliferative states in non-lung and non-prostate solid tumors (Figure 2I and 2J). However, we also observed the trend that scRNA-seq datasets from *in vitro* co-culture contained higher number of cells in proliferative states (Figure S2F). In fact, scRNA-seq data of the solid tumor-stimulated CAR-T cells with IgG hinge and CD4 transmembrane (SRP298959) was derived from *in vitro* co-culture experiments [39]. To validate this trend, we compared the ratio of cells in the proliferative states across experimental models. As expected, the percentage of cells in proliferative state was significantly lower in *in vivo* models (patients and xenografts) than *in vitro* models (p=1.30e-2 by two-sided T test) (Figure 2L). Furthermore, within *in vivo* models, CD4^+^ CAR-T cells infused into patients were less proliferative than those from animal models (p=7.54e-3 by two-sided T test). In contrast, no significant difference in percentage of proliferative cells was observed between blood cancer and solid tumor (p=7.11e-2 by two-sided T test). It has been recently reported that CAR-T cell proliferation was inhibited by the presence of M2-polarized macrophages [40]. To assess the relationship between CD4^+^ CAR-T cell proliferation and M2 macrophages, we obtained the transcriptome profiles of macrophages from the collected scRNA-seq datasets (Figure S2H), and calculated M2/M1 polarization ratio in the individual macrophages by enrichment of M1 and M2 gene signatures [41]. Expectedly, the frequency of proliferative states of CD4^+^ CAR-T cells was negatively correlated with M2/M1 ratio (Spearman correlation = 0.758, p=1.59e-2) (Figure S2I). Taken together, these results indicated that the proliferative state of CD4^+^ CAR-T cells is affected by experimental models, and *in vivo* studies are important to assess CAR-T cell proliferation during treatment.

### Suppression of NF-κB signaling in solid tumor-stimulated CD4^+^ CAR-T cell

Given the limited differences in subpopulation composition between certain solid tumors and blood cancers (Figure 2H, bottom two quadrants), we next asked how the anti-tumor activity was suppressed in these solid tumors. To this end, we performed differential expression analysis between solid tumors and blood cancers in each common cluster (Figure 2M). We observed significant down-regulation of genes related to immune cell activation, migration, and granule synthesis in the majority of the common clusters. Furthermore, solid tumor-stimulated CD4^+^ CAR-T cells down-regulated NF-κB signaling, which plays an essential role in proliferation, persistence, and survival of CAR-T cells via 4-1BB costimulation and main predictor for inflammation of immune system [42, 43]. Overall, the suppression of these immune activation pathways was a major characteristic of solid tumor-stimulated CD4^+^ CAR-T cells.

### Identification of blood cancer-specific subpopulations in CD8^+^ CAR-T cells

CD8^+^ T cell harbors cytotoxic function against cancer cells and is a prominent effector cell type in the anti-tumor immunity. CD8^+^ cells were transcriptionally segregated into two groups: a cluster unique to CAR-induced datasets, and a cluster common to both CAR- and non-induced datasets (Figure 3A). Similarly to CD4^+^ cells, high expression of cell cycle-related genes was also observed in the cluster unique to CAR-induced datasets (“CD3^+^ unique”), suggesting that CAR induction enhances proliferation of T cells (Figure 3B and 3C). Unlike CD4^+^ cells, *CD3* family genes were holistically expressed in CD8 cluster, representing CD8^+^ T cells (Figure S3A). Only a small population of *CD14* and *CD86-*expressing cells, involved in antiviral responses [44, 45], showed cytokine production (Figure 3D).

**Figure 3.**
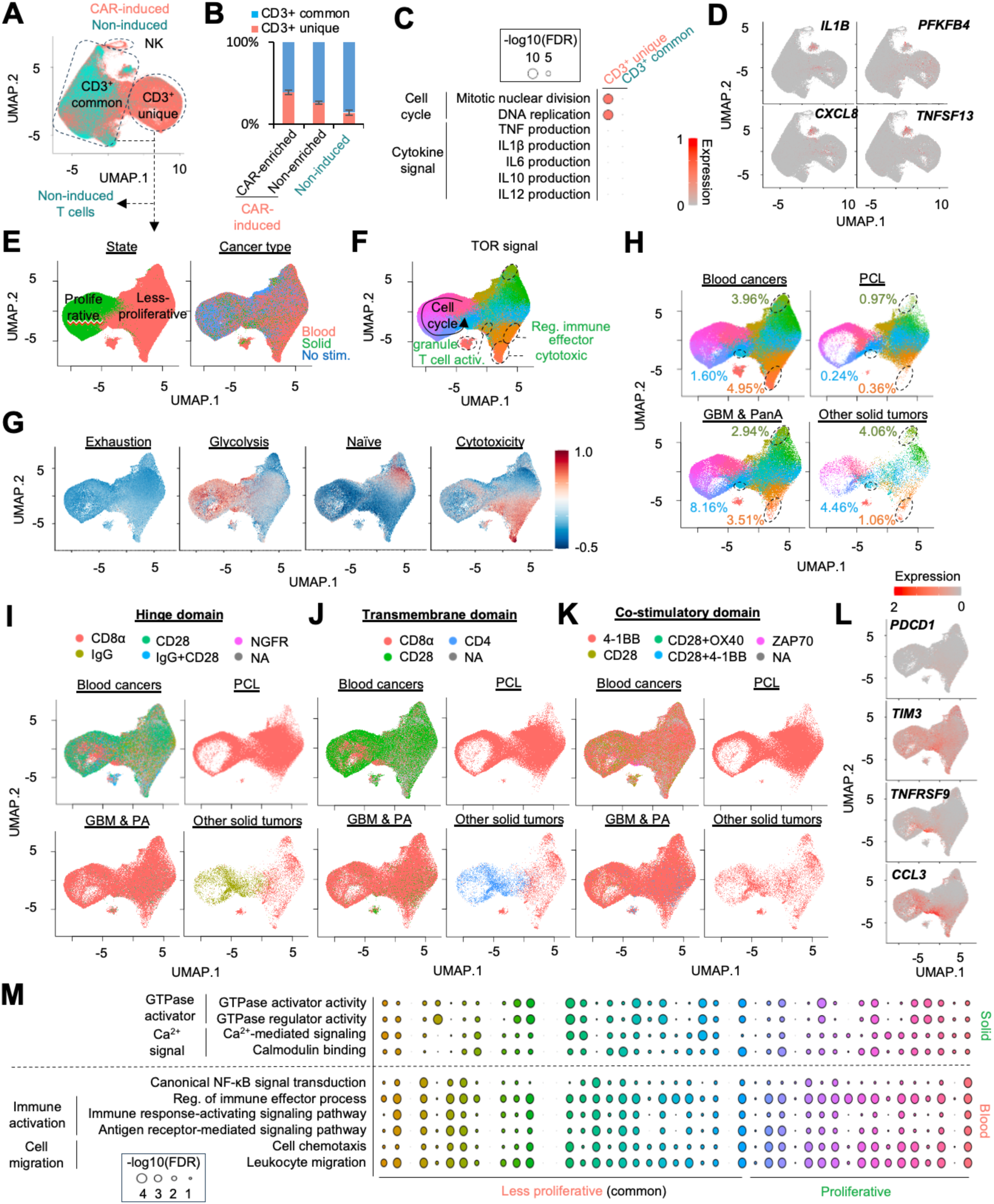
Identification of unique characteristics in blood and solid tumor-stimulated CD8^+^ CAR-T cells. **A.** UMAP plot of individual CD8^+^ T cells from 40 different projects. CAR- and non-induced T cells were distinguished by different colors. **B.** Enrichment of CD8^+^ T cell fraction in CAR- and non-induced T cells **C.** Representative significant GO terms between CAR- and non-induced CD8^+^ T cells. **D.** Feature plot representing expression of cytokine release syndrome-related genes in CD8^+^ T cells. **E.** UMAP plot of individual CD8^+^ CAR-T cells colored by proliferative states and cancer stimulation. **F.** UMAP plot of individual CD8^+^ CAR-T cells colored by clusters. Representative annotations are also shown. **G.** Feature plot representing enrichment of CD8^+^ T cell gene signatures. **H.** UMAP plot of individual CD8^+^ CAR-T cells distinguished by target cancer types. Colors correspond to clusters in Figure 3E. **I-K.** UMAP plot of individual CD8^+^ CAR-T cells distinguished by types of (**H**) hinge, (**I**) transmembrane, and (**J**) co-stimulatory domain. **L.** Feature plot representing expression of *PDCD1*, *TIM3*, *TNFRSF9*, and *CCL3*. **M.** Representative GO terms for differentially expressed genes between blood and solid tumor-stimulated CD8^+^ CAR-T cells in each common cluster. Circle size represent -log10(FDR).

We identified a few small clusters that were predominantly detected in one project (“SRP475734”) which performed scRNA-seq of CAR-T cells with and without co-injection of NK cells [30] (Figure S3C). Notably, more than 98.6% of cells in these clusters were derived from scRNA-seq libraries with NK cell co-injection. Therefore, these small clusters correspond to the co-injected NK cells and were excluded from subsequent analyses.

The re-normalization of CAR-T cell clusters revealed that certain less-proliferative clusters were unique to blood cancer-stimulated CD8^+^ CAR-T cells (Figure 3E). For example, clusters representing cytotoxicity were predominantly enriched in blood cancer-stimulated CD8^+^ CAR-T cells (Figure 3F, 3G and S3E). Clusters related to granule production, regulation of immune effector process, and T cell activation were also enriched in blood cancer-stimulated CD8^+^ CAR-T cells (>80%). Since these functions are essential for the antitumor activity of CD8^+^ T cells, scarce production of effector proteins may be a potential cause of CAR-T cell limited efficacy against solid tumors.

### Derivation of distinct CD8^+^ CAR-T subpopulations in PCL stimulation

PCL is the most aggressive form of MM and is characterized by the presence of circulating plasma cells [46]. Although cytotoxic CD8^+^ CAR-T subpopulations were substantially detectable in the majority of blood cancer types (Figure S3F), PCL stimulation did not result in sizeable production of a cytotoxic CD8^+^ CAR-T cell population (Figure 3H). In contrast, CAR-T cells against typical MM with the same CAR structure (CD8α hinge/transmembrane and 4-1BB co-stimulatory domain) could generate substantial amount of the cytotoxic subpopulations (Figure S3G) [32], suggesting that the difference in CD8^+^ CAR-T cell composition between PCL and MM may not be related to CAR construct. In addition to the cytotoxic subpopulation, PCL stimulation also infrequently generated CD8^+^ CAR-T subpopulations characterized by mTOR signaling (Figure 3F), which negatively regulate anti-tumor activity of CAR-T cells [47]. Since the PCL scRNA-seq data were obtained from a patient with complete remission [46], the scarcity of these subpopulations may actually assist the potent anti-tumor activity of CD8^+^ CAR-T cells for PCL.

### Derivation of exhaustion-like CD8^+^ CAR-T subpopulations in solid tumors and in certain blood cancers

In addition to blood cancers, the distinct composition of CD8^+^ CAR-T subpopulations was also observed across solid tumor types (Figure S3H). Among solid tumor types, cytotoxic subpopulations make up a relatively high percentage of all CD8^+^ CAR-T cells in GBM and pancreatic adenocarcinoma (Figure 3H), though recent clinical trials against these solid tumors displayed limited efficacy due to the exhaustion of CAR-T cells [10, 48]. We found that a specific CD8^+^ CAR-T cell subpopulation was predominantly derived from solid tumor datasets rather than blood cancer datasets (blue in Figure 3H, S3F-S3H). Notably, several T cell exhaustion markers, such as *TIM3* (*HAVCR2*), *TIGIT*, and *CTLA4*, were highly expressed in this cluster, although a typical marker, *PDCD1* (*PD-1*), was not enriched [25] (Figure 3L and S3I). This cluster was characterized by high expression of *TNFRSF9* and *CCL3,* which were other exhaustion-related genes [49-52]. The development of PD-1-independent exhaustion in T cells under solid tumor stimulation is concordant with a recent clinical trial which showed that anti-PD-1 treatment hardly improves the efficacy of CAR-T cell therapy against solid tumor [10]. Therefore, the blockage of alternative exhaustion pathways may be a potential strategy for the improvement of CAR-T cell therapy efficacy.

Although the TNFRSF9^+^/CCL3^+^ exhaustion-like CD8^+^ CAR-T cell subpopulation was barely detectable in blood cancer datasets, this subset was present in some B-ALL-stimulated CD8^+^ CAR-T cells (Figure S3F). In addition, the occurrence of this cluster seemed to vary depending on the CAR construct (Figure 3I and 3J). By comparing the percentage of CD8^+^ CAR-T cells within this cluster across hinge and transmembrane domain types in B-ALL, we found that CAR-T cells with CD8α hinge and transmembrane domains were overrepresented (Figure S3J). This observation implies that CAR design is an important factor to restrict CAR-T cell exhaustion.

As observed in CD4^+^ CAR-T cells, CD8^+^ CAR-T cells also significantly down-regulated the genes involved in NF-κB signaling, immune cell activation, and migration in solid tumors (Figure 3M). Furthermore, several CD8^+^ CAR-T cell clusters significantly elevated gene expression associated with GTPase activator activity. RASA2 a GTPase-activating protein and was newly identified as an intracellular checkpoint of T cell signaling by CRISPR screening [53]. We found that *RASA2* expression was significantly elevated in various CD8^+^ CAR-T cell clusters in the solid tumor datasets (Figure S3K). These observations could help the exploration of alternative checkpoint blockage strategies for the adaptation of CAR-T cells for solid tumors.

### Distinct CD8^+^ CAR-T cell composition across experimental models and patients

Like CD4^+^ CAR-T cells, the ratio of the proliferative states was significantly reduced in *in vivo*-based experimental models (patients and xenograft) (Figure S3L). Furthermore, we found that the ratio of certain CD8^+^ CAR-T subpopulation also varied across patients, in particular, between responders and non-responders. A cluster characterized by granule secretion was substantially detected in patients with relapse, but limited in those with durable remission (Figure S3M). Interestingly, this cluster highly expressed alarmins (*S100A8* and *S100A9*) that negatively impact on CAR-T cell anti-tumor activity (Figure S3N) [54]. In addition, a recent study demonstrated that *S100A8* and *S100A9* in T cells contributed to the disruption of innate immune response in severe COVID-19 patients [55]. Overall, these alarmin-expressing cells are a potential biomarker of response to CAR-T cell therapy.

### Inferring blood cancer and solid tumor effects on CAR-T cells by deep learning

Given the unique subpopulations and molecular characteristics of blood cancer- and solid tumor-stimulated CD4^+^ and CD8^+^ CAR-T cells, we next asked whether these characteristics can be automatically detected by a machine learning approach. Our group previously developed a computational tool, scIDST, that can infer heterogeneous cellular characteristics from sample annotations using a deep learning model with weak supervision framework [56]. For example, CAR induction level is not uniform across CAR-T cells and was previously reported to be positively associated with their exhausted phenotypes [57]. Although alignment of scRNA-seq reads on CAR construct is a straightforward approach, its number was too small (∼1 or 2 reads) to accurately measure CAR induction levels. To address CAR induction of individual T cells, we trained the weakly supervised deep learning model from the mixture of CAR- and non-induced scRNA-seq libraries (Figure 4A), and inferred CAR induction levels (Figure 4B) (See Methods). Notably, previously defined gene signatures of CD4^+^ and CD8^+^ CAR^high^ T cell were significantly increased along the inferred CAR induction levels (Figure 4C) [57]. Furthermore, T cells with high inferred CAR levels were predominantly detected in the solid tumor-specific CD4^+^ clusters and TNFRSF9^+^/CCL3^+^ exhaustion-like CD8^+^ clusters (Figure 4D). These results were consistent with a previous report showing that CAR^high^ T cells are more vulnerable to exhaustion [57]. Similarly, we hypothesized that the effects of blood cancer and solid tumor stimulation are also not unique across individual CAR-T cells, and inferred them with the weakly-supervised deep learning model. Interestingly, the inferred effects of blood cancer and solid tumor stimulation were significantly higher in the blood cancer- and solid tumor-enriched subpopulations, such as cytotoxic CD8^+^, TNFRSF9^+^/CCL3^+^ exhaustion-like CD8^+^, and TGFβ/SMAD and glutamate signal-related CD4^+^ CAR-T cell clusters, than in common clusters (Figure 4E-F). Overall, these results suggested that the deep learning model can recognize the distinct molecular features of CAR-T cells between blood cancer and solid tumor stimulation.

**Figure 4.**
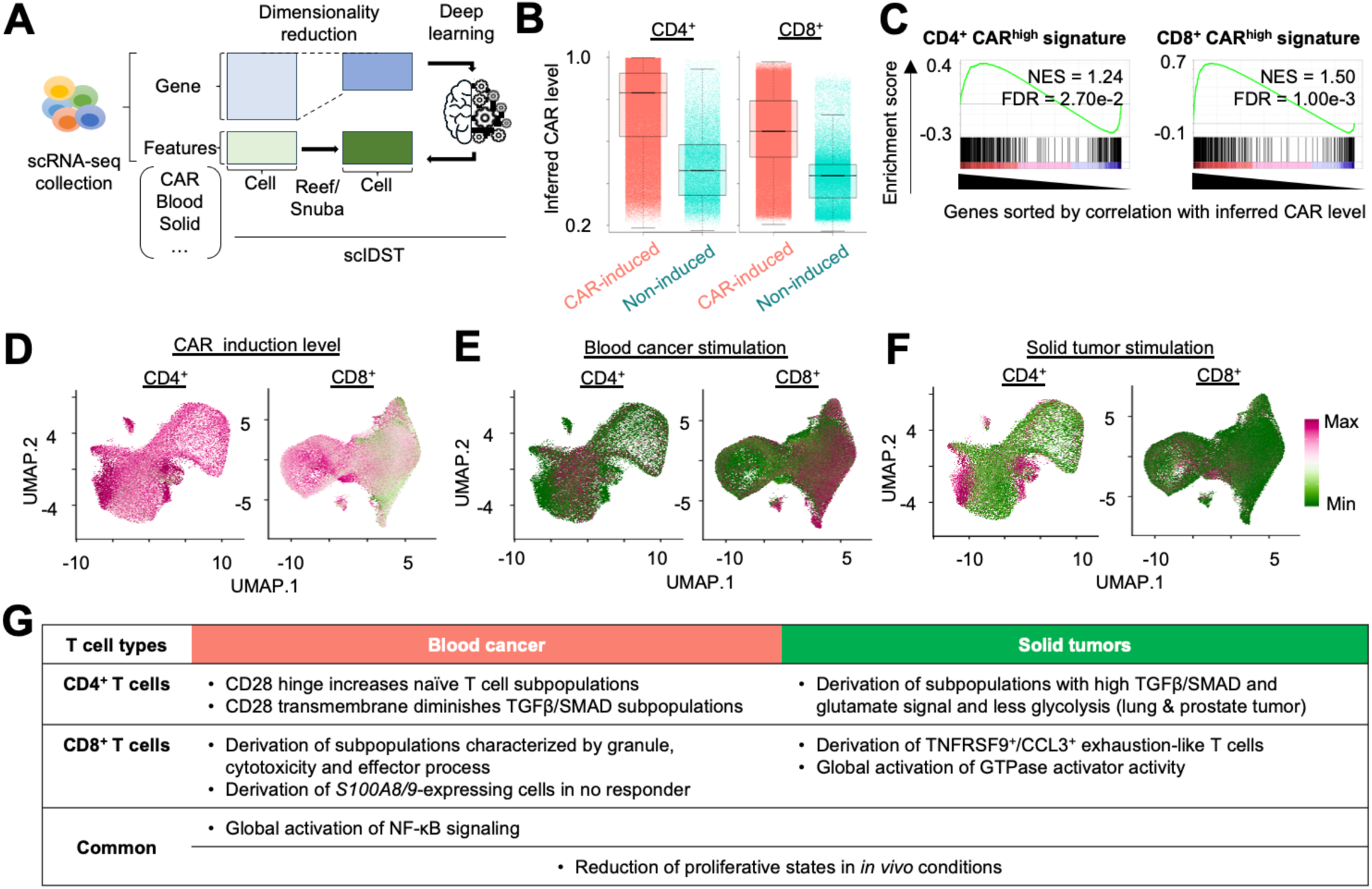
Inferring effects of blood cancer and solid tumor stimulation by deep learning. **A.** Schematics of deep learning training for inference and assessment of CAR induction level, blood cancer and solid tumor effects. **B.** Inferred CAR levels in individual CD4^+^ and CD8^+^ T cells in CAR- and non-induced datasets. **C.** Enrichment of CD4^+^ and CD8^+^ CAR^high^ T cell gene signature in CAR-T cells with high inferred CAR level. **D-F.** UMAP plots of individual CD4^+^ and CD8^+^ CAR-T cells colored by (**D**) CAR induction level and effect of (**E**) blood cancer and (**F**) solid tumor stimulation. **G.** Summary of molecular characteristics of CD4^+^ and CD8^+^ CAR-T cells between blood cancer and solid tumor stimulation

To further assess the trained deep learning model, we also inferred the effects of the blood cancer and the solid tumor stimulations on individual CAR-T cells in additional scRNA-seq datasets (Figure S4A, Table S2). Similarly, CAR-T cells were classified into less proliferative and proliferative states (Figure S4B and S4C). We observed that a portion of less proliferative clusters displayed high inferred solid tumor effects in both CD4^+^ and CD8^+^ CAR-T cells, and was characterized as high TGFβ/SMAD/glutamate signals and *TNFRSF9* and *CCL3* expression, respectively (Figure S4D-S4G). Although a small portion of CD8^+^ CAR-T cells showed high inferred blood cancer effects, these cells highly expressed cytotoxic gene signatures (Figure S4H). Taken together, the weakly-supervised deep learning model is a robust approach to predict the existence of blood cancer- and solid tumor-enriched CAR-T cell subpopulations.

## Discussion

The synthetic analysis of multiple scRNA-seq datasets has tremendous benefits for assessing robustness and reproducibility of the biological discoveries [58, 59]. Recent atlas-scale studies have revealed unique molecular characteristics in durable responders and non-responders to CAR-T cell therapy [60-62]. However, these studies focused on specific blood cancers (e.g. ALL, AML) or CAR constructs, and the differential molecular responses between blood and solid tumors and even across blood cancers were not fully investigated. In this study, using comprehensive collection of scRNA-seq, we found distinct subpopulation derivation and signaling induction between blood cancer- and solid tumor-stimulated CAR-T cells (Figure 4G). Solid tumor stimulation preferentially generated CD4^+^ CAR-T cell subpopulations characterized by the suppression of antitumor immunity, whereas cytotoxic and immune effector CD8^+^ CAR-T cell subpopulations are more predominantly generated under blood cancer stimulation. Solid tumor stimulation also preferentially generated TNFRSF9^+^/CCL3^+^ exhaustion-like CD8^+^ subpopulations. As a common characteristic in CD4^+^ and CD8^+^ CAR-T cells, NF-κB signaling was globally suppressed by solid tumor stimulation. We also found that the subpopulation derivation was also different across hinge and transmembrane domain types. In particular, the proportion of immunosuppressive CD4^+^ CAR-T and TNFRSF9^+^/CCL3^+^ exhaustion-like subpopulations was higher in CD8α-hinge and transmembrane domain than CD28. Furthermore, the subpopulation composition was also affected by experimental models; proliferation of CD4+ CAR-T cells is decreased in *in vivo* conditions, probably due to the existence of anti-tumor ineffective macrophages.

The emergence of CAR-T cell therapy has invigorated the movement of personalized immunotherapy for irremediable diseases. Its technologies are continuously improved and also expanded into the treatment of non-cancerous diseases [63]. We believe that our comprehensive collection of scRNA-seq data is a precious resource to tackle the obstacles facing CAR-T cell therapies against solid tumors and improve their clinical efficacy.

## Materials and Methods

### Data collection rules and preprocessing

To minimize technical biases, we collected scRNA-seq libraries derived from 10X Genomics Chromium, which is the most widely used single-cell platform. SRA or BAM format files were downloaded from NCBI Short Read Archive (SRA) database, and converted into FASTQ files by fastq-dump function in SRAtoolkit (v2.10.8) or bamtofastq software (v1.3.2) from 10X Genomics, respectively (Figure S1A). Some FASTQ files were directly downloaded from SRA or through Amazon Web Services. The scRNA-seq data collection used in the manuscript were retrieved by keyword search with “CAR-T cell” or other related terms on March 22^nd^, 2024 (1^st^ collection). We did not include any scRNA-seq libraries for which the access to raw data is restricted. One scRNA-seq library (SRX21787289) was removed from subsequent analyses because of missing raw data (lack of R1 file). Another RNA-seq library (SRX5310382) was also removed due to its very low read depth (<5M reads). The target cancer types, experimental models (patients, xenograft, or *in vitro*), and CAR constructs were manually collected from literature (Table S1). Unclear information (e.g. detail was not provided) was labeled as “NA”. All raw sequence data were then aligned to human GRCh38 reference genomes by CellRanger (v7.0.0) with default parameters. Since the libraries of SRP420341 project were single-cell multiome (RNA+ATAC) sequencing, we used CellRanger-arc (v1.0.1) for the alignment. Only the count matrices for RNA were used for subsequent analyses.

Quality control and batch effect correction were performed by Seurat package (v5.0.3) in R (v4.3.1) [58, 64]. Briefly, among over 3 million cells, we first filtered out approximately 0.9 million low-quality cells that had more than 10% mitochondrial reads, <2.75 or >3.85 of log10(nCount_RNA+1), and <3 or >4.25 of log10(nFeature_RNA+1) (Figure S1B). Subsequently, 255,672 cells without expression of an immune cell marker, *CD45* (*PTPRC*), were excluded as non-immune cells. The remaining immune cells were further divided into CD4^+^, CD8^+^, double positive T cells, and other immune cells by existence of scRNA-seq reads in *CD4*, *CD8A*, *CD8B*, and *CD8B2*. For example, CD8^+^ T cells had scRNA-seq reads in either *CD8* variants (*CD8A*, *CD8B*, and *CD8B2*), but not in *CD4*. The other immune cells did not have any scRNA-seq reads in *CD4*, *CD8A*, *CD8B*, and *CD8B2*. We also excluded genes that were detected in less than 10 cells or had less than 10 reads from subsequent analyses (Figure S1C). The feature-barcode (gene-cell) matrices were merged by each project, and normalized by being divided by the total counts for that cell and multiplied by scaling factor (=10,000). The top 2,000 highly variable features were then identified by vst method with FindVariableFeatures function. Dimensionality reduction was also implemented by principal component analysis (PCA), and 1-30 principal components were used for subsequent reciprocal PCA-based integration by IntegrateLayers function with “k.weight=50, method = CCAIntegration” options. After data integration, individual cells were projected into two-dimensional space by UMAP. Cell clusters were identified by a shared nearest neighbor modularity optimization with resolution=2. The potential droplets were predicted by scDblFinder (v1.16.0) with default parameters [65] (Figure S1E).

Identification of differentially expressed genes between CAR- and non-induced T cells was implemented by FindMarkers function with test.use=“wilcox” option. Genes as more than 4 fold change and less than 0.05 adjusted p-value were defined as differentially expressed genes. Gene Ontology (GO) analysis was then conducted on the differentially-expressed genes using GOstats Bioconductor packages (v2.68.0) [66]. GO terms with less than 0.05 false discovery rate were defined as statistically significant (Figure 2C and 3C).

### The re-normalization of CAR-T cell clusters

CAR-T cell clusters were subset from total CD4^+^ and CD8^+^ T cell data, and re-normalized by PCA and IntegrateLayers with “k.weight=30, method = CCAIntegration” options. One dataset, SRP489891, was excluded from the re-normalization analyses due to its strong bias to central memory T cells [21] (Figure S2C). In addition, in CD8^+^ T cells, we excluded clusters corresponding to NK cells (Figure S3C). Several datasets, whose total number of CAR-T cells were less than 30, were also excluded. The clusters were identified by a shared nearest neighbor modularity optimization with resolution=2, and grouped into proliferative and less proliferative clusters by expression patterns of cell cycle genes (*TOP2A*, *MKI67*, *PCNA*, and *MCM2*) (Figure S2D and Figure S3D). The cluster markers were identified by FindAllMarkers function, and defined by more than 2 fold change and less than 0.05 adjusted p-value. Overrepresented GO terms in each cluster were analyzed by GOstats as described above (Figure S2E and S3E). The cluster composition was compared by target cancer types, experimental models, and CAR constructs (hinge, transmembrane, and co-stimulatory domain) (Figure 2I-K and 3I-K). Since almost all CARs employed scFv variable fragment and CD3ζ activation domain, we excluded them from the comparative analysis. Differentially expressed genes between blood cancer- and solid tumor-stimulated CAR-T cells were identified by FindMarkers function in each cluster. GO enrichment analysis was performed by GOstats as described above (Figure 2M and 3M).

### Gene signature analysis for T cell functions

Curated gene sets for CD4^+^ and CD8^+^ T cell functions were obtained from Pan-cancer T cell atlas [25]. Enrichment of these T cell function-related gene set was assessed in CD4^+^ and CD8^+^ CAR-T cells by AddModuleScore function in Seurat (Figure 2G and 3G).

### Macrophage polarization analysis

Macrophage clusters were identified from the other collected immune cells by expression of its markers (e.g. *LYZ*) (Figure S2H). The enrichment of M1 and M2 gene signatures, which were collected from the literature [41], was assessed by AddModuleScore function in Seurat. M2/M1 ratio was calculated dividing M2 module score by M1 module score in individual cells and averaged in each project. Spearman correlation between M2/M1 ratio and the ratio of proliferative states and its statistical significance were calculated by cor.test function in R with method=”spearman” function (Figure S2I).

### Analysis of CD14^+^CD86^+^CD8^+^ T cell cluster

Average log-normalized read count was calculated in the cluster expressing *CD14* and *CD86*, and then subtracted by that of the other clusters. Gene Set Enrichment Analysis (GSEA) was implemented to genes sorted by ratio using GSEA software (v4.1.0) with C5.GO.BP.v2024 gene set database, 100 permutations and nocollapse of gene sets [67] (Figure S3B).

### Deep learning inference

The python scripts for deep learning and weak supervision were downloaded from GitHub repository (https://github.com/ytanaka-bio/scIDST) [56]. Briefly, the count matrices for CD4^+^ and CD8^+^ T cells were generated from Seurat objects (count matrices after quality control). 25,000 cells were randomly selected from the count matrices of CD4^+^ and CD8^+^ T cells, and used for the training of an autoencoder that allows us to do dimensional reduction in multiple independent datasets (*autoencoder.py* with “-x 50” option). The count matrices of CD4^+^ and CD8^+^ T cells were then dimensionally reduced by the trained autoencoder (*autoencoder.py* with “-r 0” option) and used for subsequent weak supervision and classification steps. All features were labeled as binary (e.g. 1=*in vivo*, 0=*in vitro*) and converted into probabilistic labels by Reef/Snuba automating weak supervision (*reef_analysis.py* with default parameters) [68]. Finally, a multi-layered artificial neural network was trained by the dimensionally reduced matrices and the probabilistic labels (*classifier_analysis.py* with train function and “-x 50” option).

As test datasets for the trained deep learning model, additional 31 scRNA-seq libraries from 3 different projects were further retrieved on October 7^th^, 2024 (2^nd^ collection) and downloaded from NCBI SRA (Table S2). Raw sequence data and quality control were preprocessed in the same way as described in the “Data collection rules and preprocessing” section with CellRanger and Seurat respectively. The deep learning models trained by version 1 datasets were used to infer CAR induction levels and the effects of blood cancer and solid tumor by *classifier_analysis.py* with predict function and default parameters.

### Quantification and Statistical Analysis

Statistical details and software used for various types of data analyses in this work are cited in the appropriate sections in the STAR Methods. Briefly, we used t.test() function to perform two-sided t test for the identification of differentially expressed genes, p.adjust() for the multiple test correction, cor.test() for the calculation of Pearson correlation and the corresponding p-value, and hyperGTest() for the hypergeometric test in GO analysis in R software. Permutation test was performed in Gene Set Enrichment Analysis with GSEA software.

### Data access and code availability

The raw UMI count matrices combining all described single-cell transcriptome profiles and the pre-trained deep learning models are available at our Mendeley repository (https://data.mendeley.com/preview/tfc4czg6sx?a=1a13acaf-4115-4bf3-bed4-571caa4f2aac). A CAR-T cell atlas (https://blood-cart-solid.cells.ucsc.edu/) is available throughout UCSC CellBrowser as an interactive web application to explore the composited scRNA-seq datasets [69]. R codes for the data preprocessing were available throughout GitHub (https://github.com/ytanaka-bio/CART_atlas/). The accession numbers of public datasets used in this study are listed in Tables S1 and S2.

## Acknowledgement

We thank Dr. Sylvie Lesage, Jean-Sébastien Delisle, Christopher E Rudd (Centre de Recherche de l’Hôpital Maisonneuve-Rosemont, Canada), and Ryuma Tanaka (Nationwide Children’s Hospital, USA) for their constructive advice for this project. This research was enabled in part by support provided by Calcul Québec (https://www.calculquebec.ca/en/) and Digital Research Alliance of Canada (https://alliancecan.ca/en).Y.T. was supported by laboratory start-up funds from Centre de Recherche de l’Hôpital Maisonneuve-Rosemont and Université de Montréal, Junior 1 & 2 Research Scholarship from Fonds de recherche du Québec–Santé (FRQS) (Dossier no. 285285 & 347381), bridge funding of Canadian Institutes of Health Research (CIHR; OGB-185739) and a transition award from Cole Foundation.

## Author contributions

Y.T. conceived of and supervised the study. Y.T. collected and preprocessed single-cell transcriptome data. Y.T. and Y.H.L. organized meta data information. E.F. and Y.H.L. analyzed single-cell datasets of CD4^+^ and CD8^+^ T cells, respectively.

## Competing interest

Y.T. works as a consultant in Colossal Biosciences. The remaining authors have no conflict of interest to declare.

## Supplemental Figures

### Table of Contents

Table S1. List of scRNA-seq datasets and sample information (version 1)

Table S2. List of scRNA-seq datasets and sample information (version 2)

**Figure S1.**
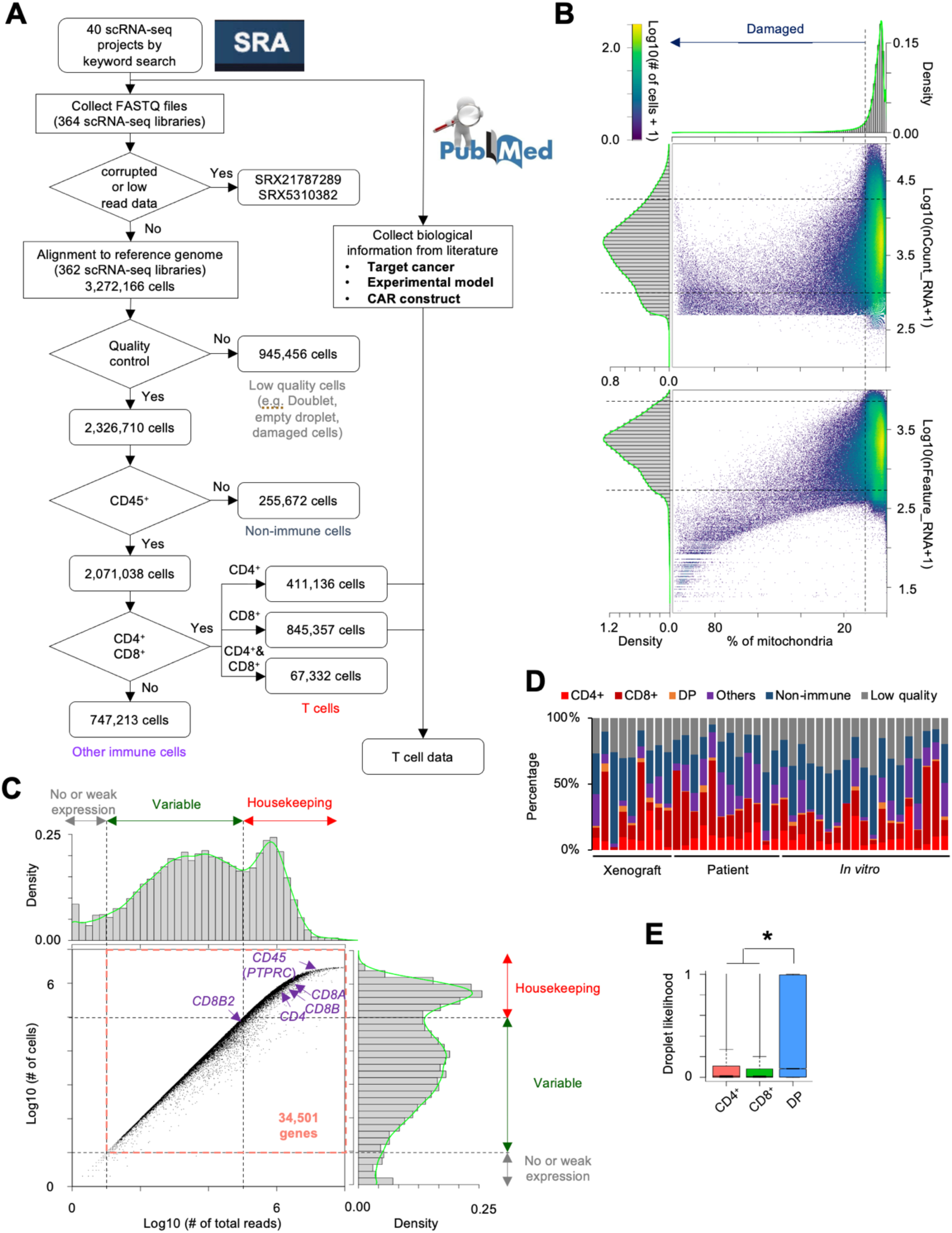
Data collection and preprocessing of single-cell transcriptomics of CAR-T cells related to Figure 1. **A.** Workflow of data preprocessing of single-cell transcriptome data for CAR-T cells **B.** Assessment of percentage of mitochondrial reads, nCount_RNA and nFeature_RNA. Dashed lines represent cutoffs for quality control. **C.** Histogram of total reads and number of detected cells in each gene. Genes with more than 10 reads and more than 10 cells were used for subsequent analyses. **D.** Percentage of each cell type and low-quality cell in each project. **E.** Comparison of inferred droplet likelihood across CD4^+^, CD8^+^, and double positive T cells. * p<0.05 by two-sided T test

**Figure S2.**
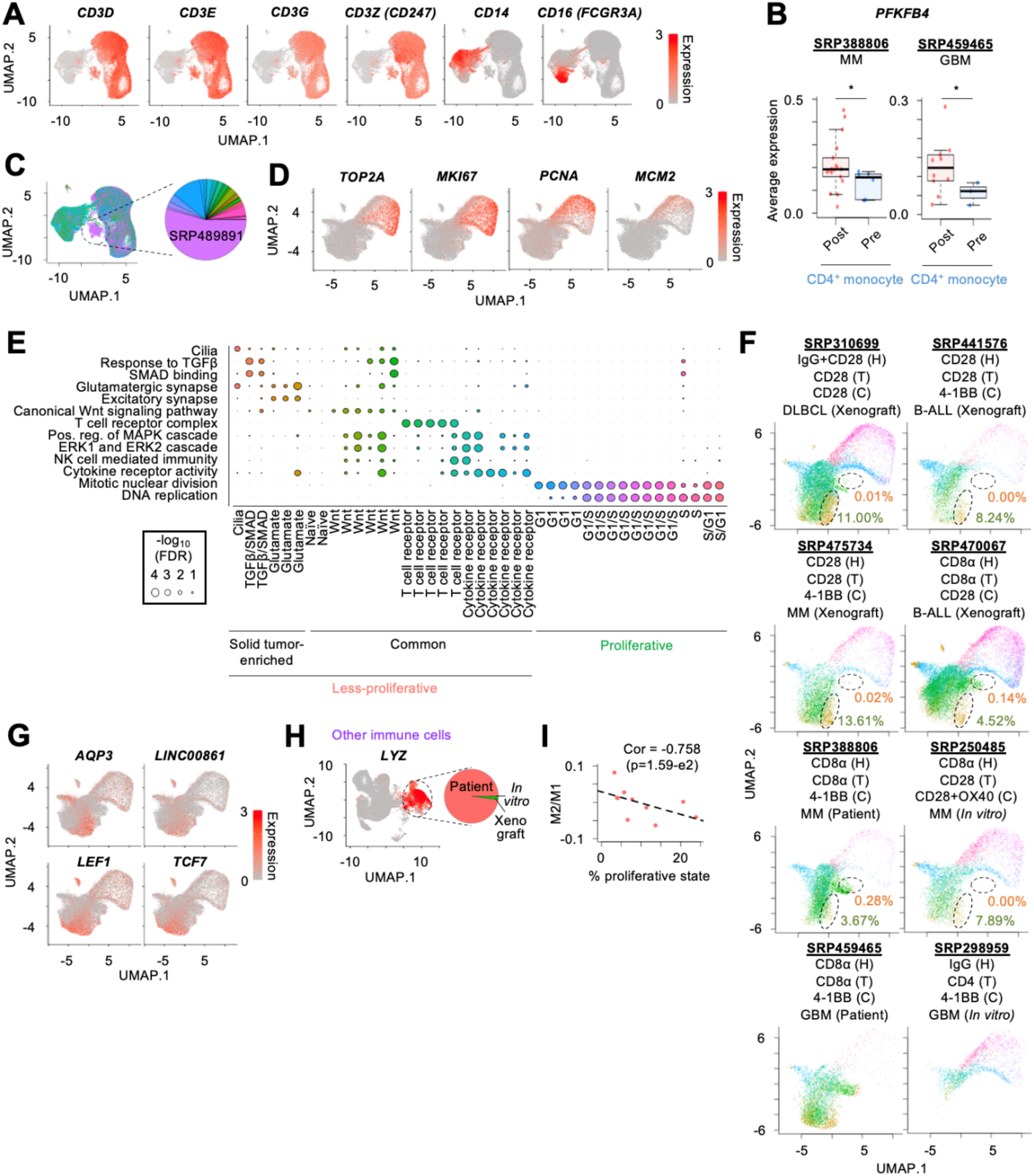
Characterizing blood and solid tumor-stimulated CD4^+^ CAR-T cells. **A.** Feature plot representing expression of *CD3* family genes and monocytic markers in CD4^+^ T cells. **B.** Comparison of average expression of *PFKFB4* in non-induced CD4^+^ T cells between post- and pre-infusion of CAR-T cells. **C.** UMAP plot of individual CD4^+^ T cells colored by project. A specific cluster is predominantly derived from one project (SRP489891). **D.** Feature plot representing expression of cell cycle genes in CD4^+^ CAR-T cells. **E.** Representative GO terms in each cluster of CD4^+^ CAR-T cells. Circle size represents - log_10_(FDR). Colors correspond to Figure 2F. **F.** Comparison of cluster composition in each representative dataset. Colors correspond to Figure 2F. Percentages of glutamate, naïve-like, and SMAD/ TGFβ clusters are also shown. **G.** Feature plot representing *AQP3* and *LINC00861* expression in CD4^+^ CAR-T cells. **H.** Feature plot representing *LYZ* expression in other immune cells **I.** Relationships between the ratio of proliferative states and average M2/M1 ratio.

**Figure S3.**
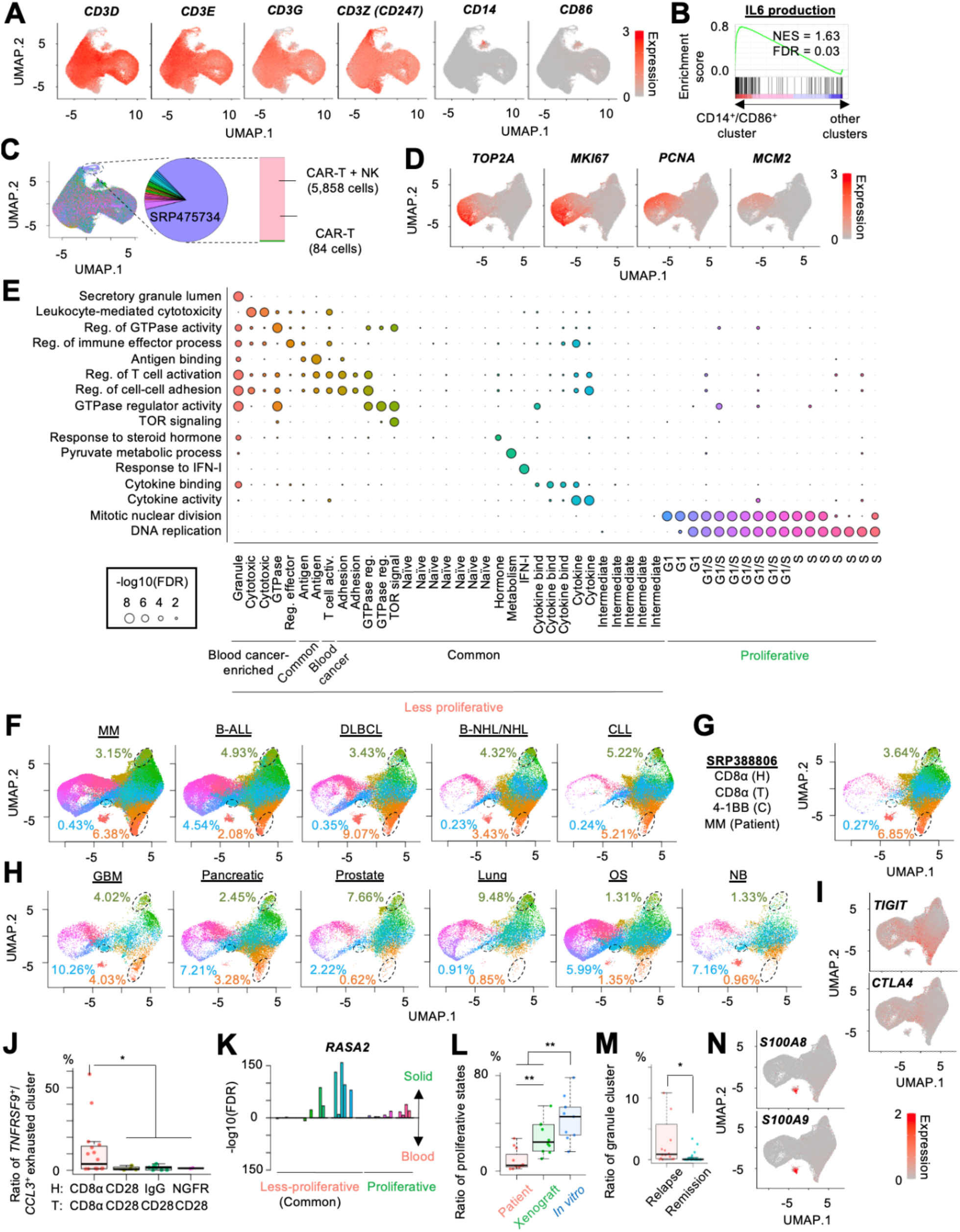
Characterizing blood and solid tumor-stimulated CD8^+^ CAR-T cells. **A.** Feature plot representing expression of *CD3* family genes, *CD14,* and *CD86* in CD8^+^ T cells. **B.** Enrichment of IL6 production-related genes in CD14^+^CD86^+^CD8^+^ T cell cluster. **C.** UMAP plot of individual CD8^+^ T cells colored by project. A specific cluster is predominantly derived from a project, SRP475734, that performed co-injection of CAR-T cells with NK cell. **D.** Feature plot representing expression of cell cycle genes in CD8^+^ CAR-T cells. **E.** Representative GO terms in each cluster of CD8^+^ CAR-T cells. Circle size represents - log_10_(FDR). Colors correspond to Figure 3F. **F-H.** Comparison of cluster composition (**F**) across blood cancer types, (**G**) in PCL, and (**H**) across solid tumor types. Colors correspond to Figure 3F. Percentages of cytotoxic, TOR signal, and TNFRSF9^+^/CCL3^+^ exhaustion-like clusters are also shown. **J.** Feature plot representing expression of *TIGIT* and *CTLA4*. **K.** Comparison of *RASA2* expression between blood cancer and solid tumor stimulation in each common less proliferative and proliferative cluster. **L.** Comparison of the ratio of proliferative states across experimental models and the target cancer types. * p<0.05 ** p<0.01 by two-sided T test **M.** Comparison of the ratio of granule cluster between relapse and durable remission samples. **N.** Feature plot representing expression of *S100A8* and *S100A9*.

**Figure S4.**
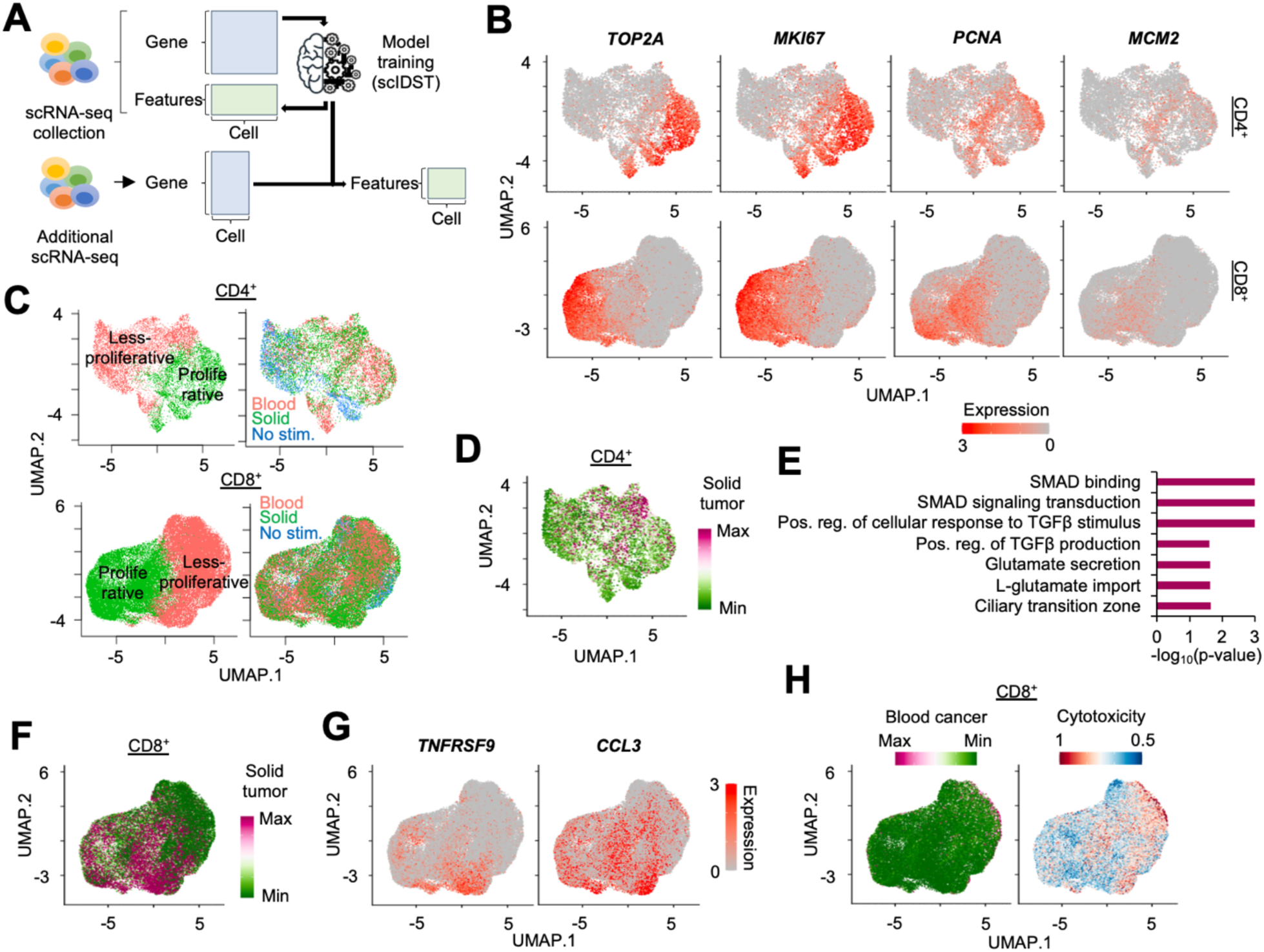
Inferring effects of blood cancer- and solid tumor stimulation on individual CD4^+^ and CD8^+^ CAR-T cells. **A.** Schematics of deep learning prediction of blood cancer and solid tumor effects. The deep learning is trained by scRNA-seq collection (362 libraries & 40 projects) and predicts the effects in additional scRNA-seq datasets (31 libraries & 3 projects). **B.** Feature plot of cell cycle genes in individual (top) CD4^+^ and (bottom) CD8^+^ CAR-T cells in scRNA-seq data collection version 2. **C.** UMAP plot of individual CD4^+^ and CD8^+^ CAR-T cells colored by proliferative states and cancer stimulation. **D.** UMAP plot of individual CD4^+^ CAR-T cells colored by effect of solid tumor stimulation. **E.** Representative GO terms for highly expressed genes in high solid tumor effects. **F.** UMAP plot of individual CD8^+^ CAR-T cells colored by effect of solid tumor stimulation. **G.** Feature plot representing expression of *TNFRSF9* and *CCL3* in CD8^+^ CAR-T cells. **H.** UMAP plot of individual CD8^+^ CAR-T cells colored by effect of blood cancer stimulation and enrichment of CD8^+^ T cell cytotoxicity gene signatures.

